# Spatial memory training reverses GirK channels modulation in the transgenic APP_Sw,Ind_ Alzheimer’s disease mouse model

**DOI:** 10.1101/2022.05.20.492780

**Authors:** Sara Temprano-Carazo, Ana Contreras, Carlos A. Saura, Juan Navarro-López, Lydia Jiménez-Díaz

**Affiliations:** Neurophysiology & Behavior Lab, Centro Regional de Investigaciones Biomédicas (CRIB), School of Medicine of Ciudad Real, University of Castilla-La Mancha, Ciudad Real, Spain; Institut de Neurociències, Department de Bioquímica i Biología Molecular, Universitat Autònoma de Barcelona, Barcelona, Spain; Centro de Investigación Biomédica en Red Enfermedades Neurodegenerativas (CIBERNED), Spain

**Author notes:** Correspondence should be addressed to Juan D. Navarro-López at or to Lydia Jiménez-Díaz at. Contributed equally (Sara Temprano-Carazo, Ana Contreras).

**Keywords:** Hippocampus, GirK, Alzheimer’s disease, APP_Sw,Ind_, spatial memory, excitatory/inhibitory imbalance

## Abstract

Alzheimer’s disease (AD) is a dementia characterized by progressive memory decline and neurodegeneration caused by the accumulation of amyloid-β (Aβ) peptides. Last findings point to an imbalance between excitatory and inhibitory neurotransmission as the initial impairment in early stages, and the hippocampus as one of the most susceptible brain areas.

The G-protein-gated inwardly rectifying potassium (GirK) channel has been proposed as a potential target to restore excitatory/inhibitory balance in amyloidosis models. Moreover, cognitive training may counteract early AD symptoms, although its effect on GirK channels remains unknown.

Here, the effect of genotype, age, and training in a hippocampal-dependent memory task on the protein expression of GirK subunits and modulators were studied using APP_Sw,Ind_ mice. Results showed a reduction of GirK2 expression as well as an increased expression of SNX27 in the hippocampus of 6-month-old APP_Sw,Ind_ mice. Training in a memory task restored GirK2 and SNX27 levels. Thus, the effect of Aβ on GirK2 could account for the excitatory/inhibitory imbalance transmission found in AD models, and training in a cognitive hippocampal-dependent task reverses this effect and lessens early Aβ-dependent AD deficits.

**Summary statement:** Aβ decreases hippocampal GirK2 expression in APP_Sw,Ind_ mice, which could contribute to early hyperexcitability found in Alzheimer’s disease models. Training in spatial memory tasks has shown to counteract this reduction.

## 1. Introduction

Over 55 million people worldwide have been diagnosed with dementia, of which 60-70% suffer from Alzheimer’s disease (AD) (Gauthier et al., 2021). AD is characterized by extracellular accumulation of β-amyloid (Aβ) plaques and intracellular neurofibrillary tangles of hyperphosphorylated *tau*, which lead to a progressive neurodegeneration and memory loss. Despite the extended research in the topic, no current therapeutic disease-modifying intervention exists (for a review, see (Jeremic et al., 2021a)).

Amyloid precursor protein (APP) transgenic mice show pathological changes resembling those of AD patients (Mucke et al., 2000). Biochemical and histological studies have shown that 2-month-old APP_Sw,Ind_ transgenic mice do not exhibit Aβ in the hippocampus, which begins to be detectable at 6 months of age, and by 12-18 month-old mice show senile plaques. Moreover, these transgenic mice show spatial memory deficits at 6 months in hippocampal-dependent memory tasks, such as Morris water maze (MWM) (Parra-Damas et al., 2014; Saura et al., 2015). Therefore, the progression of biochemical, histological and behavioral changes over time make APP_Sw,Ind_ mice an excellent murine model to compare early and late disease stages.

Hippocampal dysfunction is one of the first alterations in early stages of AD, such as excitatory/inhibitory imbalance, synaptic plasticity disruption and memory deficits (Mucke and Selkoe, 2012; Jeremic et al., 2021a). Given the proven neuronal hyperactivity (Hector and Brouillette, 2020), G-protein-gated inwardly rectifying potassium (GirK) channels have emerged as potential targets to regulate neuronal activity, based on their inhibitory role (Dascal, 1997; Djebari et al., 2021; Jeremic et al., 2021b). GirK channels are a family of K^+^ channels activated by a variety of G-protein-coupled receptors, such as GABA, dopamine, serotonin or adenosine (Lüscher and Slesinger, 2010; Slesinger and Wickman, 2015; Jeremic et al., 2021b). They form tetrameric units, assembled by a combination of their subunits GirK1-GirK4. In the brain, GirK1, GirK2 and GirK3 are widely expressed throughout different areas, while GirK4 subunit is classically related to the heart and its neuronal expression is limited (Karschin et al., 1996). Moreover, GirK2 seems to be necessary for the correct function of these channels, and GirK1/GirK2 heteromers have been identified as the main GirK channels in neurons (Pravetoni and Wickman, 2008; Jeremic et al., 2021b).

Additionally, GirK function and location depends on the effect of different modulators. Among those, the sorting nexin 27 (SNX27) and the regulator of G-protein signaling 7 (RSG7) are of special interest. SNX27 is a cytoplasmatic protein required for endosomal recycling of many important transmembrane receptors, that binds to neuronal GirK channels controlling its trafficking and surface expression (Lüscher and Slesinger, 2010; Balana et al., 2011). Furthermore, SNX27 downregulation is linked to AD through increased intracellular production of Aβ (Chandra et al., 2021). On the other hand, RSG7 functions as a GTPase activating protein (GAP) that accelerates G-protein inactivation. Accordingly, mice lacking RGS7 show profoundly slow GIRK channel deactivation kinetics (Ostrovskaya et al., 2018), disruption of inhibitory forms of synaptic plasticity and deficits in learning and memory (Ostrovskaya et al., 2014).

Previous data from our group showed that *in vitro* acute incubation with Aβ induces depolarization of CA3 hippocampal pyramidal neurons by a loss-of-function of GirK channels (Nava-Mesa et al., 2013) and deregulation GirK subunits genes (specifically, genes encoding GirK2, GirK3 and Girk4) (Mayordomo-Cava et al., 2015). Furthermore, increasing GirK activity restores hippocampal Long-Term Potentiation (LTP) and memory deficits induces by Aβ in an *in vivo* mouse model of AD (Sánchez-Rodríguez et al., 2017; Sánchez-Rodríguez et al., 2019b). However, all this promising data have been obtained in an intracerebroventricular murine model of early AD, when Aβ is soluble rather than accumulated. A recent study has provided evidence of GirK redistribution from the plasma membrane to intracellular sites and pre- and post-synaptic reduction of GIRK2 channels in two transgenic AD models, P301S and APP/PS1 (Alfaro-Ruiz et al., 2021). Nevertheless, whether GirK subunits levels are altered in the brain of mice developing AD pathological changes during aging is unknown.

Furthermore, it is well stablished that periodic cognitive training on a hippocampal-dependent memory task may mitigate memory deficits observed in AD, both in rodents (Parra-Damas et al., 2014; Martinez-Coria et al., 2015; Rai et al., 2020) and humans (Hall et al., 2009; Oveisgharan et al., 2020), as well as attenuate Aβ deposition and enhance adult hippocampal neurogenesis (Zhao et al., 2022). Accordingly, training in a spatial memory task might be able to reverse the alterations in GirK expression caused by Aβ-overexpression, with the subsequent recovery of hippocampal impairment.

Thus, the aim of the present study was to determine the effect of genotype, age and training on the protein expression of GirK subunits and its modulators, in the hippocampus of APP_Sw,Ind_ transgenic mice.

## 2. Results

The experimental design of this study is detailed in Fig. 1A. Firstly, we examined the protein levels of GirK channel subunits and modulators in the hippocampus of adult APP_Sw,Ind_ transgenic mice at early (6 months) and late (12-18 months) pathological stages (Parra-Damas et al., 2014). As shown in Fig. 1B, APP_Sw,Ind_ mice showed intracellular Aβ accumulation in the hippocampus at 6 months and amyloid plaques at 12–18 months. Thus, these ages are adequate choices to compare early and late disease stages in this murine model.

**Figure 1.**
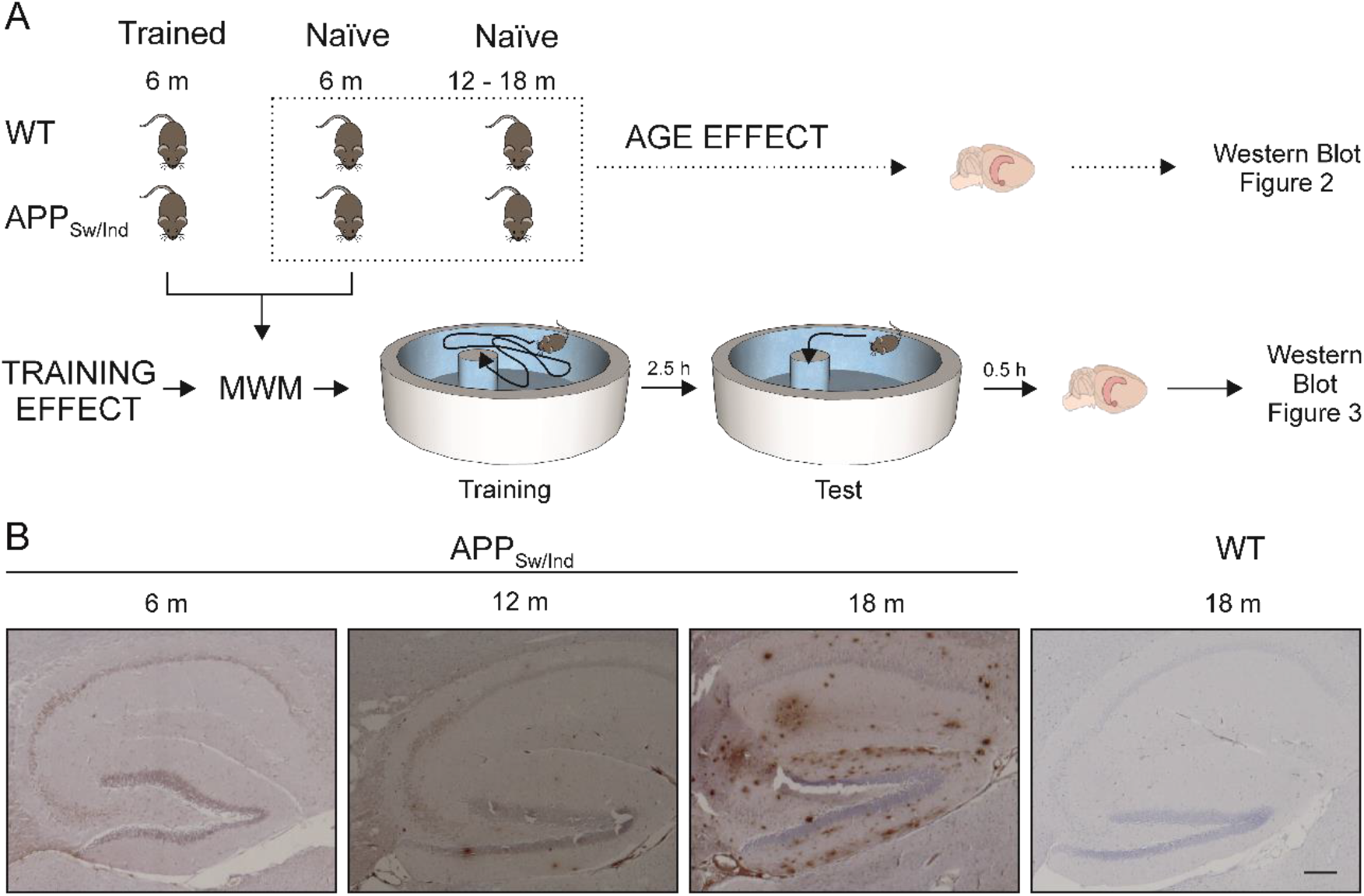
Experimental design and progression of amyloid pathology. (A) Experimental design showing number of animals for each condition, representation of training in the Morris Water Maze (MWM), hippocampal dissection for Western blot analysis, and corresponding figure numbers. (B) Age-dependent amyloid pathology in the hippocampus of APP_Sw,Ind_ mice at 6, 12 and 18 months of age. Brain sections were stained with an anti-Aβ 6E10 antibody. Scale bar: 250 μm. m, months; h, hours; WT, wild type.

Interestingly, quantitative analyses of hippocampal lysates showed a significant effect of genotype in the expression of GirK2 (F_(1,15)_ = 7.715; *p* = 0.014) and SNX27 (F_(1,14)_ = 3.974; *p* = 0.066). As shown in Fig. 2B, GirK2 was downregulated in 6 month-old APP_Sw,Ind_ mice compared to age-matched controls (t_(8)_ = 2.602; *p* = 0.0315). By contrast, the GirK modulator SNX27 was significantly upregulated in 6 month-old APP_Sw,Ind_ mice (Fig. 2F; t_(6)_ = 3.099; *p* = 0.0211) but not in control mice. These data indicate that cerebral Aβ accumulation modulates GirK channel expression.

**Figure 2.**
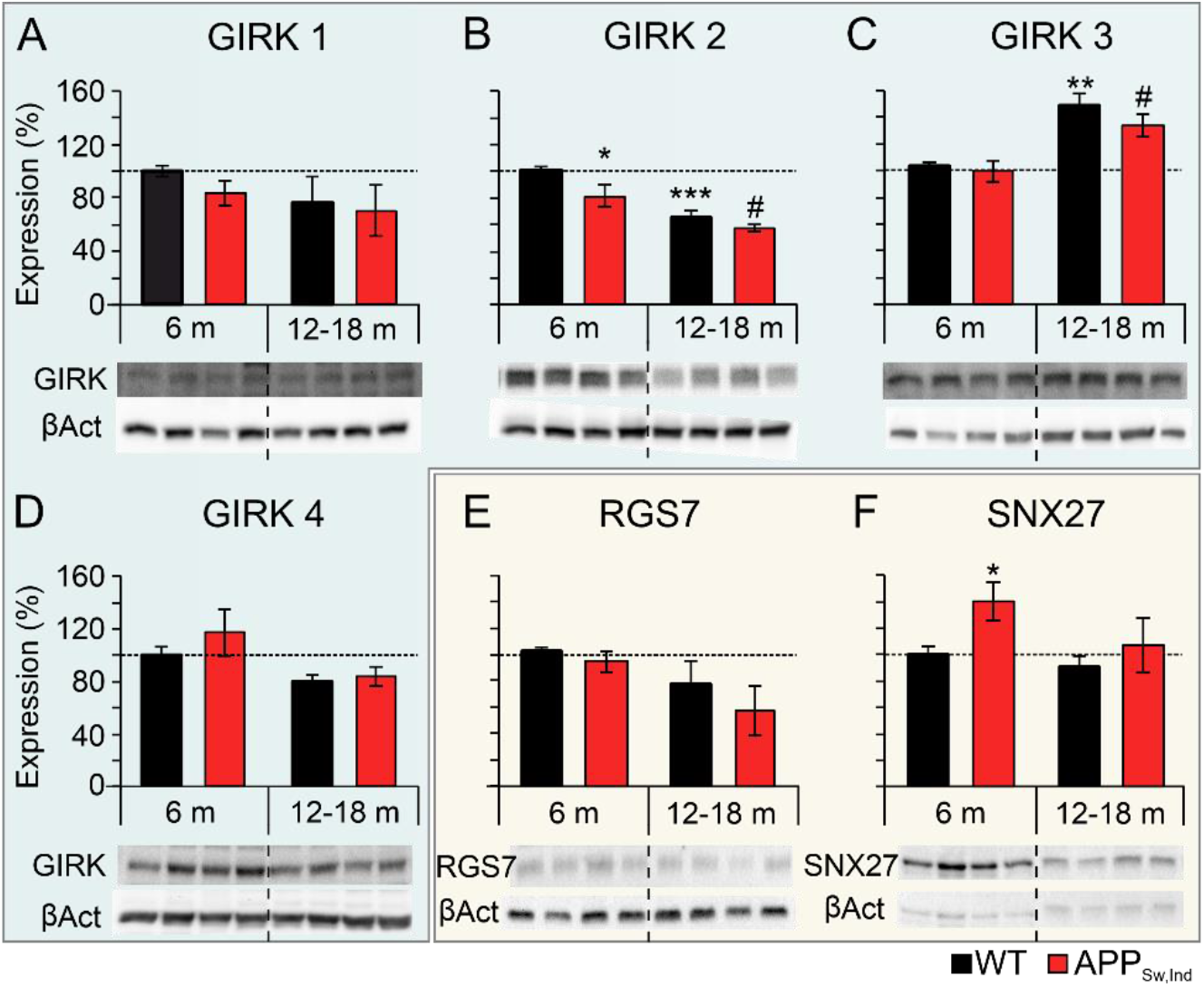
Effect of genotype and age on hippocampal levels of GirK channel subunits and modulators. Protein levels of GirK channel subunits (1-4) and modulators (RGS7 and SNX27) in the hippocampus of non-transgenic (WT) and APP_Sw,Ind_ mice at 6 and 12–18 months of age. (A) GirK1, (B) GirK2, (C) GirK3, (D) GirK4, (E) RGS7 and (F) SNX27. Data are expressed as percentage relative to 6 month-old WT mice (100%). * *p* < 0.05, ** *p* < 0.01, *** *p* < 0.001 *vs*. WT 6 month-old; # *p* < 0.05 *vs*. APP_Sw,Ind_ 6 month-old. m, months; βAct, β-actin.

Moreover, an age effect was observed in the expression of GirK2 (F_(1,15)_ = 33.973; *p* < 0.001) and GirK3 (F_(1,18)_ = 19.775; *p* < 0.001) in both WT and APP_Sw,Ind_ mice. These effects were opposite, as GirK2 expression was decreased (Fig. 2B; WT: t_(10)_ = 5.824; *p* < 0.001; APP_Sw,Ind_: t_(5)_ = 2.967; *p* = 0.0313) and GirK3 was increased (Fig. 2C; WT: t_(10)_ = 3.412; *p* = 0.0066; APP_Sw,Ind_: t_(8)_ = 2.889; *p* = 0.0202) in aged WT and APP_Sw,Ind_ mice. No statistical differences were found in hippocampal levels of GirK1, GirK4 and RGS7 in the experimental groups during aging (Fig. 2A, D and E). These data suggest that age, regardless of the genotype, varies the expression of specific GirK subunits and modulators.

Next, we analyzed the effect of spatial memory training in the MWM on the levels of GirK subunits and modulators (Fig. 1A). At 6 months of age, APP_Sw,Ind_ mice showed higher latencies to find the hidden platform and reduced number of target platform crossings and target platform occupancy in the probe trial (Parra-Damas et al., 2014). Although no significant changes were found in GirK subunits and their modulators in naïve and trained animals (Fig. 3A-F), our results showed once again a genotype effect in the expression of GirK2 (Fig. 3B; F_(1,14)_ = 4.790; *p* = 0.046) and SNX27 (Fig. 3F; F_(1,10)_ = 6.406; *p* = 0.030). However, whereas GirK2 levels were decreased and SNX27 was increased in naïve APP_Sw,Ind_ mice compared to naïve WT mice, these changes were not detected after memory training (Fig. 3B and F). These results suggest that spatial memory training was able to restore the expression of this functional subunit and its modulator in the hippocampus of APP_Sw,Ind_ mice.

**Figure 3.**
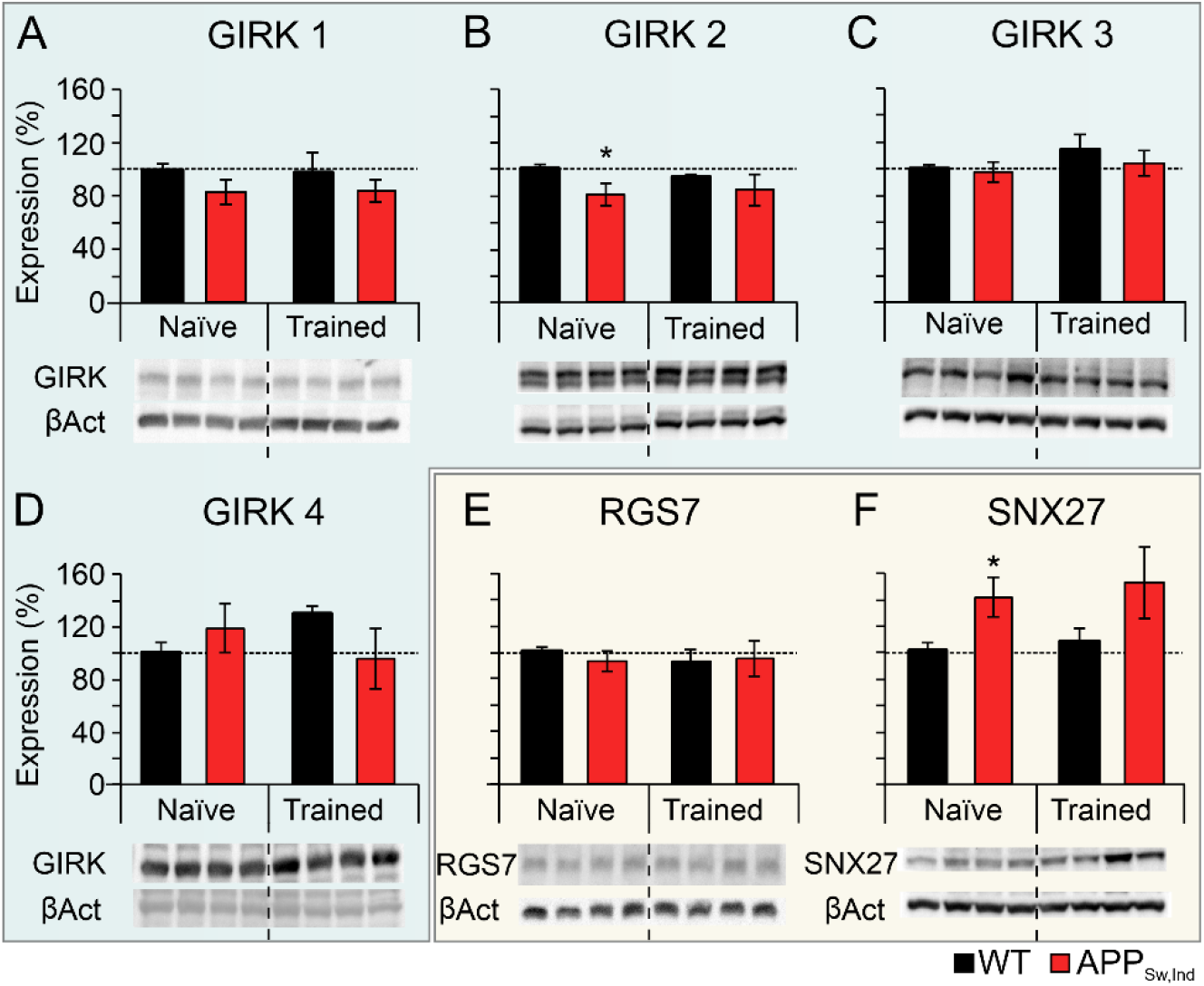
Effect of genotype and spatial memory training on hippocampal levels of GirK channel subunits and modulators. Protein levels of GirK channel subunits (1-4) and modulators (RGS7 and SNX27) in the hippocampus of naïve and memory-trained non-transgenic (WT) and APP_Sw,Ind_ mice at 6 months of age. (A) GirK1, (B) GirK2, (C) GirK3, (D) GirK4, (E) RGS7 and (F) SNX27. Data are expressed as percentage relative to untrained-naïve WT mice (100%). * *p* < 0.05 *vs*. naïve WT. βAct, β-actin.

## 3. Discussion

It has been previously demonstrated that genetic expression of GirK in the hippocampus is modulated by the biologically active fragment of amyloid-β (Aβ_25-35_) (Mayordomo-Cava et al., 2015), with deleterious effects on learning and memory processes (Mayordomo-Cava et al., 2020). The present study aimed to elucidate whether this channel is modulated in a transgenic murine model of AD. Furthermore, we studied the possible effect of age and training in a memory task on this regulation.

Young APP_Sw,Ind_ mice showed a down-regulation of the expression of GirK2 compared to WT animals. This subunit is being related to several functions, such as learning and memory, reward and motor coordination, and some pathologies like Down syndrome and epilepsy (Lüscher and Slesinger, 2010; Luján et al., 2014; Jeremic et al., 2021b). Indeed, Down syndrome patients show cerebral accumulation of Aβ and dementia symptoms during aging (Lott and Head, 2005; Moncaster et al., 2010; Cooper et al., 2012). Animals lacking GirK2 show a reduction of LTP and an increase of LTD in the hippocampus (Lüscher and Slesinger, 2010), similarly to what it is observed in AD amyloidopathy (Sánchez-Rodríguez et al., 2019a). Since this subunit is believed to be essential for the proper inhibitory function of GirK channels, its reduction could be contributing to the early hyperexcitability found in this pathology (Sanchez et al., 2012). In fact, GIRK2^-/-^ mice showed elevated motor activity in an open field and elevated lever press behavior in an operant task (Pravetoni and Wickman, 2008), which show a hyperactivity in these mice that could be induced by the underlying neural hyperexcitability. In agreement with our results, other authors have shown a reduction of GirK2 expression in the hippocampus of P301S mice, a murine model of tau pathology, yet no differences were found in APP/PS1 mice (Alfaro-Ruiz et al., 2021). This difference could be due to pathological and/or age differences between the models, since in the present study GirK2 down-expression was observed only at 6 months but not in older animals, as the ones used by Alfaro-Ruiz et al. (2021).

On the other hand, no differences in the expression of any other GirK subunit were found due to genotype. Indeed, it has already been reported that amyloidosis did not affect the expression of GirK1 (Sánchez-Rodríguez et al., 2019a; Akyuz et al., 2020). Available data regarding GirK3 is limited, however, previous work from our group showed a decreased in the genetic expression of this subunit (Mayordomo-Cava et al., 2015), while present results revealed no genotype effect. Other than the possible differences in gene and protein expression, this discrepancy could be due to differences in the experimental design, since the former was an *ex vivo* Aβ administration in hippocampal slices from 1 month-old rats (Mayordomo-Cava et al., 2015) and the latter is an *in vivo* Aβ physiological increase in older (6 and 12-18 month-old) mice.

Regarding GirK modulators, our data show an increase in the expression of SNX27 in 6 month-old APP_Sw,Ind_ mice compared to age-matched WT littermates. Contrary to this result, a recent study has linked a downregulation of SNX27 to AD, through an increased in the intracellular production of Aβ (Chandra et al., 2021). Nevertheless, this increment of SNX27 could be associated to the loss of GirK2, due to its function as a trafficking protein (Lüscher and Slesinger, 2010; Balana et al., 2011).

Moreover, our data show an age-dependent decrease in hippocampal GirK2 expression along with an overexpression of GirK3, regardless the genotype. This age-dependent change in GirK channel conformation is in line with previous studies, which showed that GirK2 progressively decreased with age, whereas GirK1 and GirK3 gradually increased during postnatal development to reach adult levels (Fernández-Alacid et al., 2011). Normal ageing is related to deficits in neurotransmission and synaptic plasticity in several brain areas (Foster and Norris, 1997; Barnes, 2003; Disterhoft et al., 2004). This translates in the impoverishment of hippocampal-dependent memory processes (Clarke et al., 2010; Warburton and Brown, 2010; Cohen and Stackman, 2015). Additionally, memory deficits in early AD have been attributed to a selective neurodegeneration in the entorhinal cortex, CA1 and the subiculum (Morrison and Hof, 2002; Giannakopoulos et al., 2009). Although CA1 seems to be the most vulnerable area of the hippocampus, there is almost no neuron loss in this region in normal ageing (West et al., 1994). Double et al. (2010) pointed to GirK2 as one possible contributor for the selective vulnerability to neural death within different brain regions. For instance, in Parkinson disease patients GirK2 is mainly expressed in the substantia nigra, where 90% of dopaminergic neurons are lost (Double et al., 2010). Thus, it seems that GirK2 expression may have a role in the vulnerability of the hippocampus to Aβ.

Interestingly, 6 month-old APP_Sw,Ind_ mice trained in the MWM did not exhibit the decrease in GirK2 that was observed in naïve transgenic animals. Plenty of studies had shown the benefits of cognitive training to slow down early AD’s symptoms (Martinez-Coria et al., 2015; Rai et al., 2020). It is relevant that a previous report indicated that, compared with trained control mice, APP_Sw,Ind_ mice showed 932 genes (88% downregulated and 12% upregulated) differentially expressed in the hippocampus. This gene profile revealed a gene cluster of 164 transcripts related to learning/memory, neurotransmission, synaptic plasticity, glutamatergic and GABAergic neurotransmission, oxidative phosphorylation and AD (Parra-Damas et al., 2014; Saura et al., 2015). Furthermore, the overexpression of GirK2 in dorsal CA1 pyramidal neurons was able to restore contextual fear learning in a GirK2^-/-^ mouse line (Marron Fernandez de Velasco et al., 2017). In this line, our results indicate that training in a spatial learning task seems to be a good strategy to compensate the GirK channel expression deficits caused by early Aβ accumulation, probably by counteracting the characteristic hippocampal hyperexcitability present in this pathology (Sánchez-Rodríguez et al., 2017; Sánchez-Rodríguez et al., 2019a; Sánchez-Rodríguez et al., 2019b). Additionally, the upregulation of SNX27 in APP_Sw,Ind_ mice was also reversed in trained mice. Together, this two results concur with the fact that SNX27 binds to neuronal GirK channels, controlling its trafficking and surface expression (Lüscher and Slesinger, 2010; Balana et al., 2011).

In conclusion, Aβ caused a decrease in GirK2 protein expression that could underlie the imbalance of excitatory/inhibitory transmission, leading to early memory deficits in AD mouse models. Moreover, training in a cognitive hippocampal-dependent task could reverse this GirK channel modulation, enhancing its inhibitory activity, and therefore lessen/ameliorating the Aβ-mediated hyperexcitability present in early AD stages.

## 4. Materials and methods

### 4.1. APP_Sw,Ind_ transgenic mice

Male APP_Sw,Ind_ transgenic mice (line J9; C57BL/6 background; *Mus musculus*), that express the human APP695 harboring the FAD-linked Swedish (K670N/M671L) and Indiana (V717F) mutations under the platelet-derived growth factor subunit B (PDGFβ) promoter, were obtained by crossing heterozygous APP_Sw,Ind_ to non-transgenic (WT) mice (Mucke et al., 2000). Age-matched male WT littermates were used as control (C57BL/6 background). Both WT and APP_Sw;Ind_ mice were genotyped individually. All experimental procedures were conducted according to the approved protocols from the Animal and Human Ethical Committee of the Universitat Autònoma de Barcelona (CEEAH 2895) and Generalitat de Catalunya (10571) following the experimental European Union guidelines and regulations (2010/63/EU).

To assess the effect of age on GirK expression, littermate WT and APP_Sw;Ind_ mice at 6 months (n = 3-8) and 12-18 months (n = 3-5) of age were used. Furthermore, to evaluate the effect of learning, 6 month-old APP_Sw;Ind_ and WT naïve mice (n = 3-8) were compared to age-matched mice trained in the Morris Water Maze (MWM) (n = 2-4), as previously described (Parra-Damas et al., 2014).

Briefly, as shown in Fig. 1A, handled mice were placed in a circular pool (90 cm diameter; 6.5 cm hidden platform) for five consecutive days (4 trials daily; 60 s per trial). Memory retention was tested 2.5 h after the last training session, and mice were sacrificed 30 min afterwards. Those times were chosen in order to get a measure of memory retention while achieving a maximum peak of gene expression, which occurs about 0.5–2 h after spatial training (Guzowski et al., 2001; Parra-Damas et al., 2014).

Animals at the proper age (6 or 12-18 month-old) or 30 min after MWM training were sacrificed by decapitation, hippocampi were dissected, and samples were frozen at −80°C until further use.

### 4.2. Immunohistochemical staining

To assess age-dependent amyloid pathology, a protocol shown to label specifically Aβ in APP transgenic mice was used, as described previously (España et al., 2010). Sagittal brain paraffin sections (5 μm) were deparaffinized in xylene, rehydrated, and incubated with 3% hydrogen peroxide. Sections were then incubated in 60% formic acid for 6 min to allow antigen retrieval, washed in 0.1 M Tris-HCl, and incubated with anti-Aβ (6E10; 1:1000; Signet, US) before immunoperoxidase staining and analysis with a Nikon Eclipse 90i microscope.

### 4.3. Western Blot

Whole hippocampal tissue samples were homogenized in ice-cold lysis RIPA-DOC buffer (Sigma, US). Protein concentration was measured using the Pierce™ BCA Protein Assay Kit (Thermo Fisher Scientific, US) according to the manufacturer’s instructions. Equal amounts of protein (10 μg) were loaded on an SDS-PAGE gel (10%) and subjected to electrophoresis. Proteins were transferred to PVDF membranes (Bio-Rad, US) using a trans-blot turbo apparatus (Bio-Rad, US). Membranes were blocked with 5% dried skimmed milk powder in Tween-TBS for 1 h. Primary antibodies (Table 1) were applied at the appropriate dilution overnight at 4 °C. After washing, appropriate secondary antibodies (Table 1) were added for 45 min at a dilution of 1/3000. Blots were detected after incubation in enhanced chemiluminescence reagent (ECL Prime; Bio-Rad, US), using the G:BOX Chemi XX6 system (Syngene, India). In order to check the equal loading of samples, blots were re-incubated with β-actin antibody as a housekeeping gene (Affinity Bioreagents, US) and data is expressed as the ratio of target protein and β-actin.

**Table 1.** Characteristics of primary and secondary antibodies used to measure protein expression by Western Blot.

| Protein | Reference | Supplier | Host | Dilution |
| --- | --- | --- | --- | --- |
| GirK1 | APC-005 | Alomone, Israel | Rabbit | 1/500 |
| GirK2 | APC-006 | Alomone, Israel | Rabbit | 1/500 |
| GirK3 | APC-038 | Alomone, Israel | Rabbit | 1/500 |
| GirK4 | APC-027 | Alomone, Israel | Rabbit | 1/500 |
| SNX27 | PA 5-23025 | Thermo Fisher Scientific, US | Rabbit | 1/100 |
| RGS7 | SC-8139 | Santa Cruz Biotech, US | Goat | 1/500 |
| β-actin | AC-15 | Sigma Aldrich, US | Mouse | 1:100000 |
| Rabbit IgG-HRP | 170-6515 | Bio-Rad, US | Goat | 1/3000 |
| Mouse IgG-HRP | 170-6516 | Bio-Rad, US | Goat | 1/3000 |
| Goat IgG-HRP | SC-2352 | Merck – Millipore, US | Monkey | 1/3000 |

### 4.4. Statistical Analysis

Two way-ANOVA was used to assess differences between genotype and age/training. When comparing only two groups, Student t test was used. Data is expressed as the mean ± S.E.M. and all analyses were performed using the IBM SPSS Statistics 24 software (SPSS Inc, US). A value of p < 0.05 was considered statistically significant. Final figures were prepared using CorelDraw v.18 Graphics Suite Software (RRID:SCR_014235).

## Author contributions

LJD, CAS and JDNL were responsible for the initial conceptualization. STC performed the formal analysis. AC was responsible for writing the original draft. AC, JDNL, CAS and LJD did the writing – review and editing. All authors have read and approved the final manuscript.

